# Biodiversity Image Quality Metadata Augments Convolutional Neural Network Classification of Fish Species

**DOI:** 10.1101/2021.01.28.428644

**Authors:** Jeremy Leipzig, Yasin Bakis, Xiaojun Wang, Mohannad Elhamod, Kelly Diamond, Wasila Dahdul, Anuj Karpatne, Murat Maga, Paula Mabee, Henry L. Bart, Jane Greenberg

## Abstract

Biodiversity image repositories are crucial sources of training data for machine learning approaches to biological research. Metadata, specifically metadata about object quality, is putatively an important prerequisite to selecting sample subsets for these experiments. This study demonstrates the importance of *image quality metadata* to a species classification experiment involving a corpus of 1935 fish specimen images which were annotated with 22 metadata quality properties. A small subset of high quality images produced an F1 accuracy of 0.41 compared to 0.35 for a taxonomically matched subset of low quality images when used by a convolutional neural network approach to species identification. Using the full corpus of images revealed that image quality differed between correctly classified and misclassified images. We found the visibility of all anatomical features was the most important quality feature for classification accuracy. We suggest biodiversity image repositories consider adopting a minimal set of image quality metadata to support future machine learning projects.

## 1. Introduction

The extensive growth in open science repositories, and, in particular, the underlying application of rich metadata has potential value for data mining, machine learning and deep learning (ML/DL). Metadata has historically been used for machine learning and automatic document classification ^1^ and there is growing attention to the role of metadata in reproducible research pipelines ^2,3^. Less common, but of paramount importance is metadata that denotes the quality of the object being represented. Metadata addressing quality control characteristics of data can support the data cleaning steps common to virtually all ML/DL analyses. In fact, computer vision is one area of particular interest where quality-specific metadata can play an important role in the selection of training, validation and test image sets. For example, Ellen et al. found the use of context metadata, consisting of hydrographic, geotemporal, and geometric data, collected or extracted from plankton images improved the accuracy of a convolutional neural network (CNN) classifier ^4^ Tang found a 7% gain in mean average precision after including GPS coordinates in a general image classification task ^5^. These studies shed light on an important area of metadata research that has broad implications for leveraging collections of digital images across nearly every scientific discipline.

One area of particular interest is specimen images, particularly given their value as a data source for species identification and morphological study ^6, 7^. The research presented in this paper, addresses this topic in the context of a National Science Foundation supported Harnessing the Data Revolution (HDR) project, Biology-Guided Neural Networks for Discovering Phenotypic Traits (BGNN). A team of information and computer scientists, biologists, and image experts are collaborating to develop a novel set of artificial neural networks (ANNs) for classifying fish species and extracting data on fish external morphological features from images of fish specimens. Unlike genomic data, specimen trait data is largely unstructured and not machine-readable. The paucity of trait data for many groups of organisms has led to efforts to apply neural network-based classification and morphological analysis to the extensive store of existing species photographic images to automatically extract trait data in an unsupervised manner. The product of these efforts, the focus of BGNN, should improve our ability to derive phenotype data from digital analogs of specimens. Metadata is recognized as an important aspect of this research, particularly in the selection of images for ML/DL.

The research presented in this paper demonstrates the value of image quality metadata for BGNN, and presents the results of baseline research on automatic specimen classification, depending on metadata quality metrics. The sections that follow provide contextual background; describe the research sample of digital images and the scheme of 22 image metadata attributes, and present the results of our baseline study. The conclusion highlights the importance of this work, specifically metadata that indicates image quality attributes important for selecting high-quality images for, in our case, biology-guided training of neural networks to extract phenotypic trait data from diverse assortments of fish specimen images.

## 2. Image Metadata Content Description and Quality

Over the last few decades, national and international support has targeted the digitization of analog specimen collections. Examples of key programs include the U.S. National Science Foundation’s (NSF) Advancing Digitization of Biodiversity Collections (ADBC) program and the European Union’s (EU) Distributed System of Scientific Collections (DiSSCo) project. Digital images produced by these initiatives have further supported the creation of large-scale specimen images repositories (e.g., The Phthiraptera Database http://phthiraptera.info/sid; Burke Museum Paleontology Database, https://www.burkemuseum.org/; Morphosource, https://www.morphosource.org/), where the images and their associated metadata are freely available for research and education purposes. The search, discovery, and use of images from these and other collections is highly dependent on the metadata associated with the image objects. A number of different metadata schemes are used across these projects, due, in part to the diversity of scope of the projects and the make-up of the project teams.

The MODAL framework ^8,9^ provides a mechanism for understanding the range and diversity of metadata standards used to describe digital specimens, as their domain foci (general to specific), including the extent of support for image quality. Metadata applicable to analog images, which is our immediate focus, falls broadly into two classes, 1.) descriptive metadata that is humanly generated by a curator; and 2.) technical metadata that is automatically generated by technology used to capture the image. The Dublin Core Metadata Standard and the Darwin Core Standard are among the most popular schemes used for describing digital specimens. These metadata standards are more descriptive covering *specimen name* and *topical* and *geo-spatial aspects*, with limited coverage of technical aspects. On the technical end, Dublin Core’s ***dc:type*** or ***dc:description*** properties or Darwin Core’s ***dwc:dynamic*** property may be used to record information impacting object quality, but these properties are still limited coverage of quality measures. The recently developed Audubon Core metadata standard for multimedia objects includes a couple of metadata properties classed under the Service Access Point Vocabulary that support some aspects of quality assessment. For example, ***Image-Height*** and ***Image-Width*** can give an indication of quality, but without knowledge of the ideal height and width it is difficult to make a clear assessment of image quality.

A richer set of metadata properties giving insight into image quality is found in the more technically oriented metadata standards identified in **Table 1**. The example metadata properties need to be measured against parameters that define image quality to be of value. The Digital Imaging and Communications in Medicine (DICOM) standard is an exception; this extensive scheme with over 200 metadata properties includes ***imageQuality*** as a metadata property, and supports scoring on a scale of 1 to 100.

**Table 1:** Example Technical and Biomedical Metadata Standards

| Metadata Standard | Primary Focus | Metadata quality property |
| --- | --- | --- |
| Preservation Metadata: Implementation Strategies | Long term preservation | Fixity |
| Exchangeable image file format (EXIF) | Image formats | X/Y dimensions, compression, color space |
| DICOM <sup>10</sup> | Medical imaging | imageQuality (1-100) |

Semantically-oriented ontologies and even controlled vocabularies, can also be used to indicate value. **Table 2** identifies two ontologies, and example semantics, that indicate image quality, and **Figure 1** illustrates the class-hierarchy where the entity ***Thing***, representing an anatomical feature and aspect of color (e.g., hue and saturation) can encode object quality.

**Table 2:** Selected Ontologies

| Ontology | Primary Focus | Semantics/metadata values |
| --- | --- | --- |
| Biomedical Image Ontology (BIM) <sup>11</sup> | Biomedical images | Image filters, ImagePreProcessing, ImagePostProcessing |
| OntoNeuroBase <sup>12</sup> | Neuro imaging | structure of interest, orientation, segmentation result |

**Figure 1:**
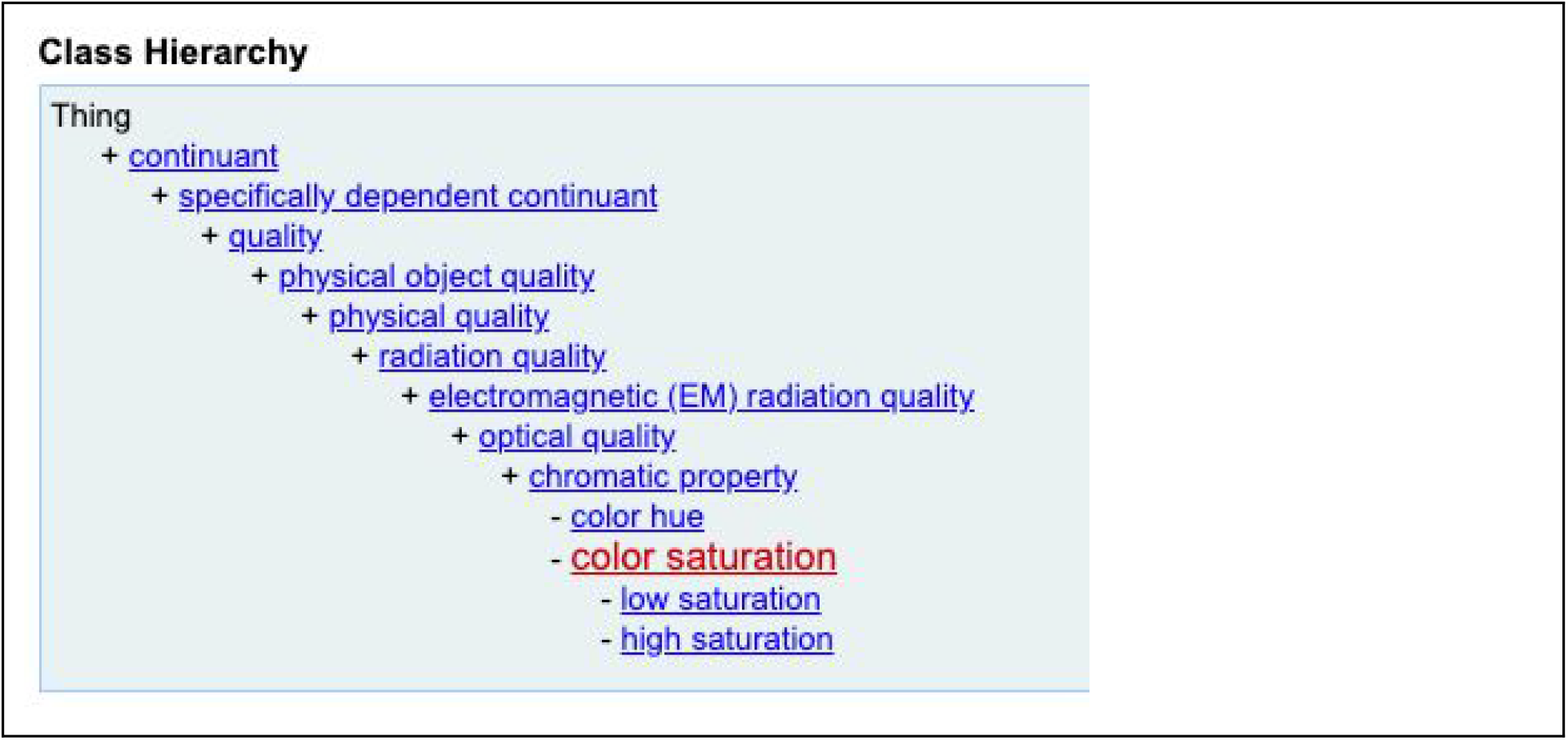
Phenotype And Trait Ontology: Example of Quality relating to color saturation and other factors.

Overall, the schemes identified here range in their focus on content description, supporting discovery with metadata properties such as a specimen’s scientific name, geographic location, provenance, and collector’s name, to technical aspects that aid in access and can help to determine aspects of quality, particularly when the parameters of what determines quality are known. Our assessment finds there does not yet exist a targeted metadata standard that captures the types of object – in our case, specimen image quality – necessary for our work in BGNN.

The need for adequate metadata to support our efforts to aggregate a sufficient quantity and variety of teleost fish images for experiments in species classification, trait segmentation and ultimately automated phenotyping in a supervised machine learning context, led us to examine a number of large curated image repositories. We initially explored using images from the iDigBio Portal (https://www.idigbio.org/portal) the national repository of NSF’s Advancing Digitization of Biodiversity Collections program, a 10-year effort to digitize data on specimens in U.S. biodiversity collections. The iDigBio Portal reportedly hosts 160,000 media files on fishes. However, searching the images, visualizing them online and obtaining copies presented challenges. A random check of roughly 5,000 images subsampled from 100,000 media files representing a number of families and genera of teleost fishes detected misidentified fish species and non-fish specimens such as plants and crustaceans. Moreover, few of the images had metadata for filtering based on quality and the metadata were incomplete in most of these instances.

We ultimately settled on using images of fish specimens from the Great Lakes Invasives Network (GLIN), one of the NSF ADBC Thematic Collections Networks. The images used in this study were obtained from the Illinois Natural History Survey Fish Collection, one of six fish collections that participated in the GLIN project.

While the metadata associated with the GLIN collections is not extensive and does not indicate image quality, the overall quality of the images was useful to serve the needs of the project, provided we gathered basic metadata to confirm the quality of individual images. Conversely image metadata may be rich and follow a standard, but that standard may not include aspects of image quality relevant to machine learning. We found image quality varied substantially, and perhaps more importantly, unevenly with respect to taxa within image repositories, which may belie both individual variation in photography and batch effects associated with submitters. This challenge, ultimately, led BGNN research to identify a set metadata properties to track image specimen quality, and the analysis demonstrating the value of these properties.

## 3. Quality Metadata for Machine Learning

The challenges we faced obtaining images of fish specimens from image repositories for use in the BGNN Project have implications across neural networks using images. The process we undertook to determine which images would be useful to us and manually annotated facets of image quality affect species classification accuracy. This manual approach contrasts with the automated image quality assessment (IQA) which is the focus of most machine learning image quality work.

IQA is an established and active area of research within computer vision. IQA research is concerned with the automated assessment of image quality for perception by both humans and machines, and the creation of robust image classification, segmentation, or detection models that are resilient to low-quality images. Reference-based IQA techniques (full-reference or FR) attempt to quantify the effects of distortions such as those imposed by compression or other forms of blur, noise, or loss of contrast. No-reference IQA algorithms have no access to high-quality reference images during inference ^13^. Both FR and NR approaches rely on ratings from several human observers to establish subjective measures for training. Databases such as the Tampere Image Database ^14^ serve as fixed gold standard references for these attempts at quantifying and modeling image quality.

Unlike IQA, domain-specific annotated quality assessment is conducted by trained annotators for the purposes of grading key relevant features. For example, in fishes the ability to see and count features like scales and ray-fins is crucial to species identification.This intense work, captured in a set of metadata properties, informed our research goals.

## 4. Research Goals

The overall aim of our research was to examine the importance of *image quality metadata* for species classification. Our goals were the following

1. Determine if the annotated image quality affected classification accuracy
2. Determine which specific quality annotations were most important for classification accuracy
3. Make recommendations for future image quality metadata in large image repositories

## 5. Methods

To examine the above goals, we conducted an empirical analysis that involved the following computational steps. Except where noted, analyses were performed using R and Python. Research data is located at https://bgnn.org/iqm reproducible source code is available at https://github.com/orgs/hdr-bgnn/iqm.

### 5.1 Sample and Evaluation Metrics

The dataset used for this study comprises 23,807 digital images of fish specimens from the Illinois Natural History Survey (INHS) Fish Collection that were produced for the Great Lakes Invasives Network Project ^15^. After checking the images for file duplications, errors with image or file formats, institution code, catalog numbers and suffixes to file names, the images were transferred to a file server from which they could be shared with other researchers in the BGNN project. Specimen collection information (occurrence records) for the images were gathered from FishNet2 ^16^ and the scientific names were updated using Eschmeyer’s Catalog of Fishes ^17^.

Based on our experience with other fish specimen image datasets available online, we defined a set of metadata properties to record image quality for the digitized fish specimens (**Table 3**). The set of properties forms a metadata scheme for capturing image quality, and is based on the expertise of members at Tulane University’s Biodiversity Research Institute, with feedback from members of the Metadata Research Center, Drexel University. Team members included informaticians, fish experts, and data entry technicians, who defined and refined the metadata scheme over a period of several months, as they processed specimen images. The scheme includes 22 metadata properties, requiring the content-value of a categorical concept, free text, a Boolean operator, or a score. A web-based form (Figure 2), and an underlying SQL-based database help to expedite capturing the metadata content. The main purpose of capturing this metadata is to derive a set of quality features (metadata properties) for filtering and retrieving digitized specimens and determining how the quality features impact image processing and machine learning.

**Table 3:** Quality Metadata Fields. The quality-annotated data set contained 19 genera and 56 species of fishes.

| Metadata: Quality property | Data type | Description |
| --- | --- | --- |
| 1. if_fish | Boolean | Whether or not fish are in the image. |
| 2. fish_number | integer | Number of fish in the image. |
| 3. if_ruler | Boolean | Whether or not a ruler in the image. |
| 4. if_label | boolean | Whether or not a label in the image. |
| 5. if_label_name_correct | boolean | Whether or not the image name matches the number on the label. |
| 6. if_label_catalog_number_correct | boolean | Whether or not the catalog number is correct. |
| 7. label_detailed | categorical | Whether or not the label is detailed and has full information. |
| 8. if_colorbar | boolean | Whether or not a colorbar is in the image. |
| 9. if_each_fish_label | boolean | Whether or not each fish specimen has a label. |
| 10. non_specimen_objects | freetext | Identify objects in image that are not fish specimens. |
| 11. if_overlapping | boolean | Whether or not objects in the image overlap or hide any parts of the fish. |
| 12. specimen_angled | categorical (1-12 based on clock face) | Number on a clock face the fish's head is pointing to. |
| 13. specimen_view | categorical | The surface(s) of the specimen visible (left side visible when head points left) |
| 14. if_curved | categorical | Whether or not the body of the fish specimen is curved or straight. |
| 15. if_missing_parts | boolean | Whether or not any parts of the fish are cropped from view. |
| 16. if_parts_visible | boolean | Whether or not all parts of the fish (including fins) are visible. |
| 17. if_fins_folded | boolean | Whether or not any of the fins are folded. |
| 18. brightness | categorical | The brightness of the fish specimen.(dark, bright, normal) |
| 19. if_background_uniform | boolean | Whether or not the background of the image is uniform. |
| 20. if_focus | boolean | Whether or not the fish specimen is in focus. |
| 21. color_issues | categorical | Identify color-related issues with the specimen (e.g. Contrast, Saturation etc.) |
| 22. image_quality | Integer (1-10) | The overall quality of the image. |

**Figure 2.**
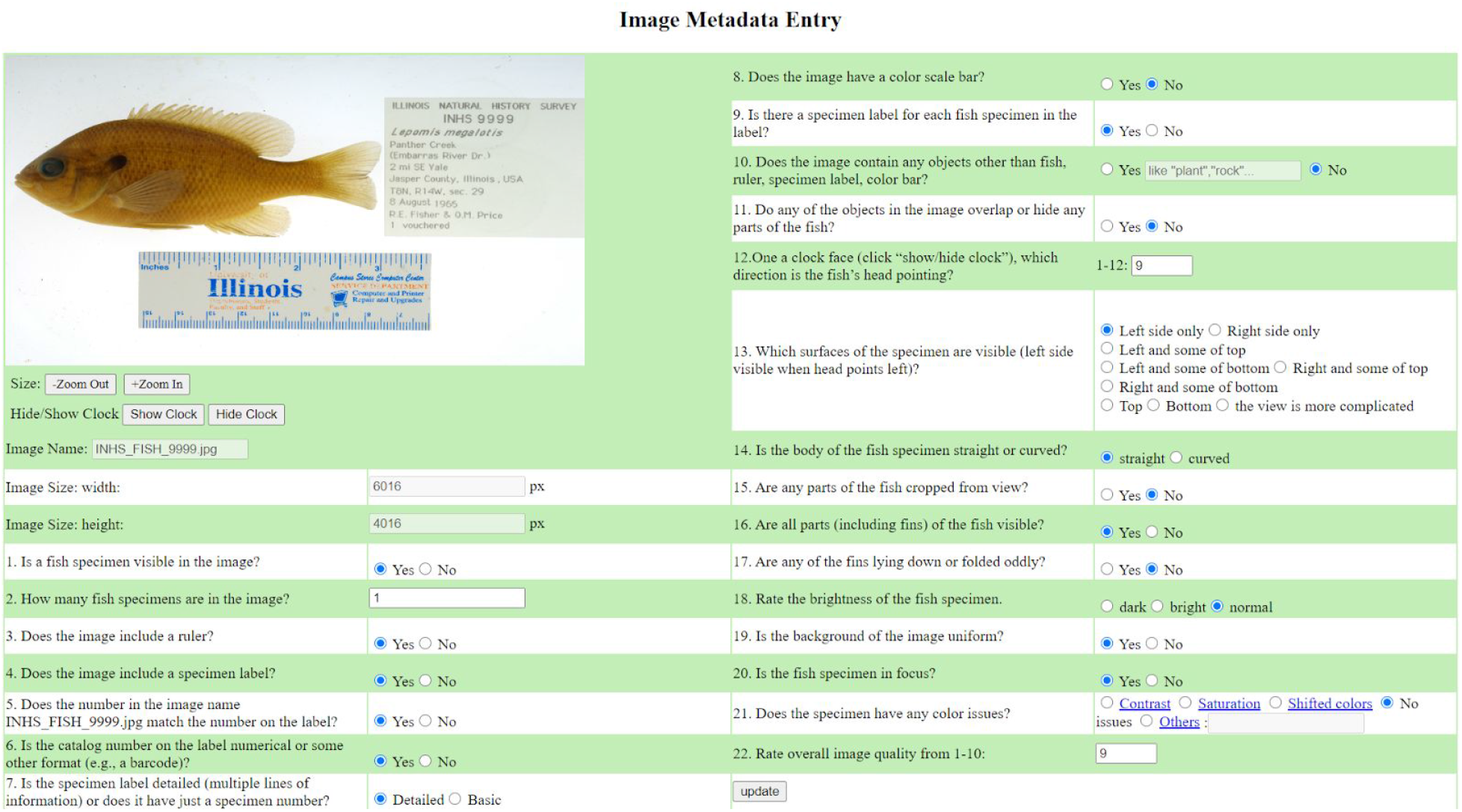
Web form for quality metadata entry

The image quality score used in this study is an integer value between 1 to 10 that represents the evaluator’s gestalt opinion of the quality of the image and its usefulness for further analysis. The evaluator assigns the score to an image after answering 21 quality-related questions about the image. However, the overall image quality score is independent of the other quality assessment questions and doesn’t integrate data from responses to any of these questions.

A score of 8-10 would be for images that are good to excellent, a score of 5-7 would be for images that have some issues but may be usable and a score of 1-4 would be for images that have major problems and are unusable.

### 5.2 Descriptive statistical analysis of quality

A basic exploratory data analysis was performed on quality metrics. Quality averages by taxonomic groups (genus and species) were examined in order to understand potential biases.

### 5.3 Implementation of a CNN-based classification pipeline

A convolutional neural network image classification pipeline was developed using PyTorch ^18^ with Torchvision ^19^ extensions. Genera (genus groups) and species (genus + specific epithet combinations) were trained and inferred simultaneously using a novel multi-classifier model, called a Hierarchy-Guided Neural Network ^20^. Several hyperparameters, including learning rate, regularization lambda, early stopping patience were tuned prior to this quality analysis.

### 5.4 Classification accuracy using high vs low-quality subsets

Using the composite median image_quality score of 8, we divided the data set into low-quality and high-quality subsets. Some species are inherently more visually similar to others, so in a classification scenario, an unbalanced distribution of taxa would confound our aim of measuring the isolated effect of image quality. To address this we sampled images based on equal distributions of individuals by species (*Esox americanus, Gambusia affinis, Lepomis gibbosus, Lepomis humilis*, *Lepomis macrochirus, Lepomis megalotis, Lepomis cyanellus, Notropis atherinoides*, *Notropis blennius, Notropis buccata, Notropis hudsonius, Notropis stramineus, Notropis volucellus*, *Noturus gyrinus*, and *Phenacobius mirabilis*) totaling 221 individuals among high and low quality subsets.

### 5.5 Quality features distinguishing correctly and incorrectly identified species

Using a dataset of 1703 quality annotated images with 20 or more individuals per species (in order to achieve enough training and test data), the holdout test set of 341 images (17 species over 8 Genera) was then divided into correctly classified and misclassified images. Quality features between these two subsets were compared, and pairs of correct/incorrect within species were examined closely.

At the time of this publication, metadata annotations indicating quality have been created for a total of 1935 images. Table 3, column 1, lists the metadata properties that form the bases of quality assessment; column 2 and 3, follow with each metadata property’s data type, and provides a brief description.

In order to prepare images for classification, Advanced Normalization Tools in R (ANTsR)^21^ was used to subtract background shading, rulers, and labels using a supervised segmentation with a training set developed in 3D Slicer ^22^. This “fish mask” was used to isolate fish images for downstream steps.

Early attempts at classification using a subset of 53 species belonging to 13 genera, with each species having 50 images, produced a mean F_1_ score (2*((precision*recall)/(precision+recall)) of 0.757 (±0.017 SD) for species classification and 0.910 for genus classification.

Our analysis focused on comparing low and high-quality images that were roughly balanced by genus and species composition, in order to control for the effect of inherent differences in identification difficulty that vary among taxa. We noted that image quality varied non-randomly among genera and species (Figure 3 & Figure 4), perhaps due to batch effects as well as anatomic differences between fish taxa that affect photographic fidelity.

**Figure 3:**
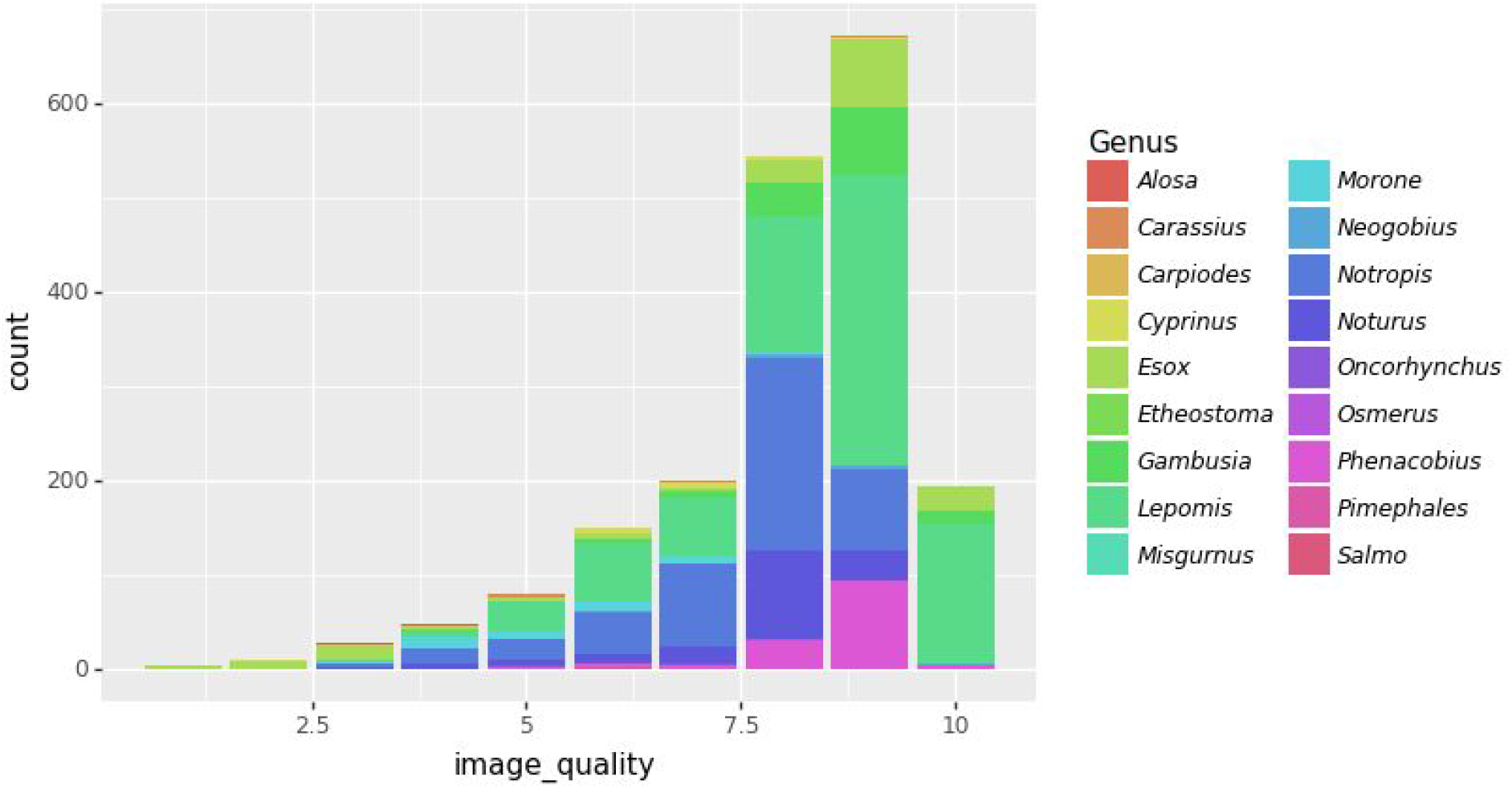
A histogram of manually annotated image quality scores across the 18 genera

**Figure 4:**
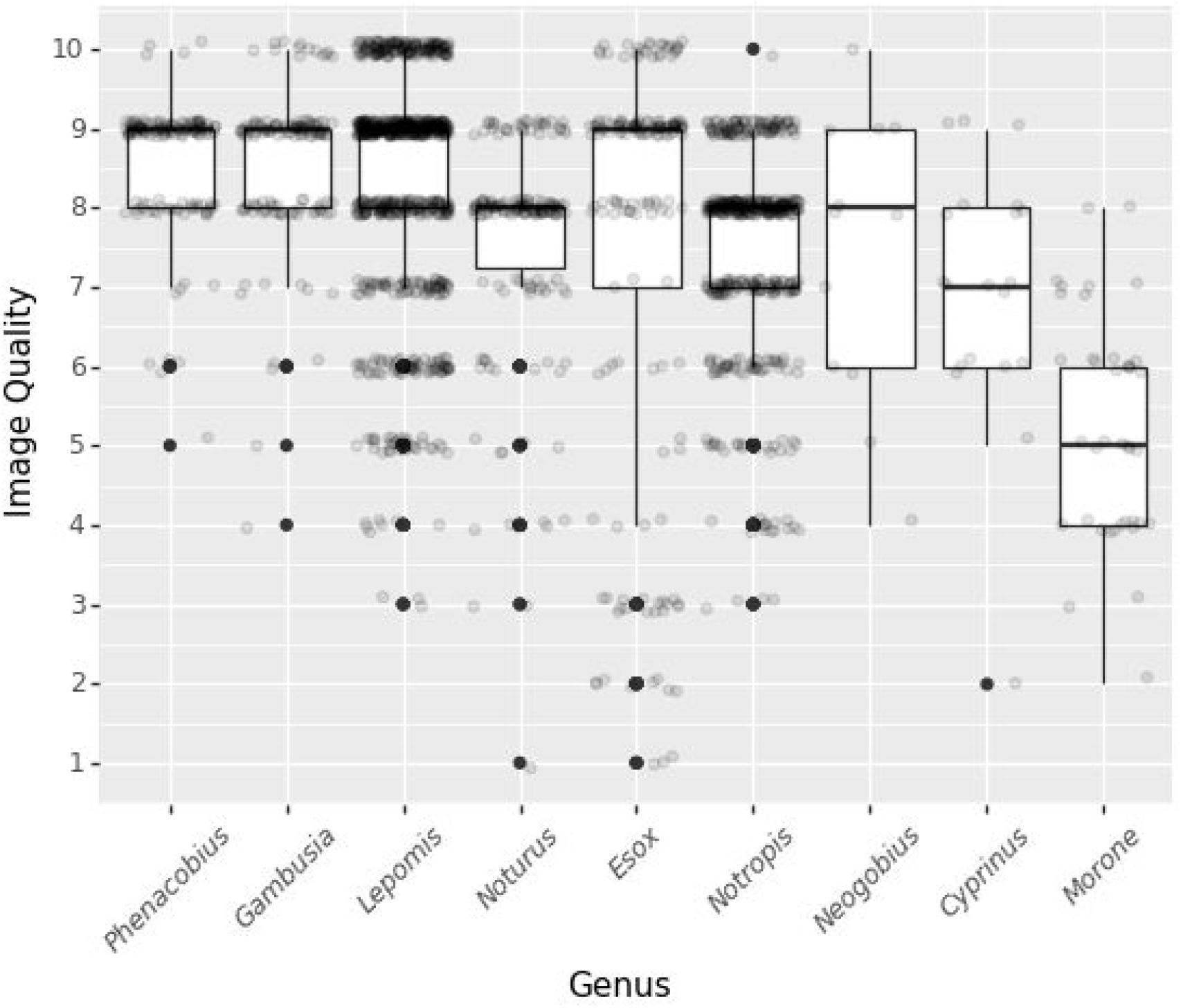
Quality scores by genera with more than 10 individuals clearly shows variability within and between genera

**Figure 5:**
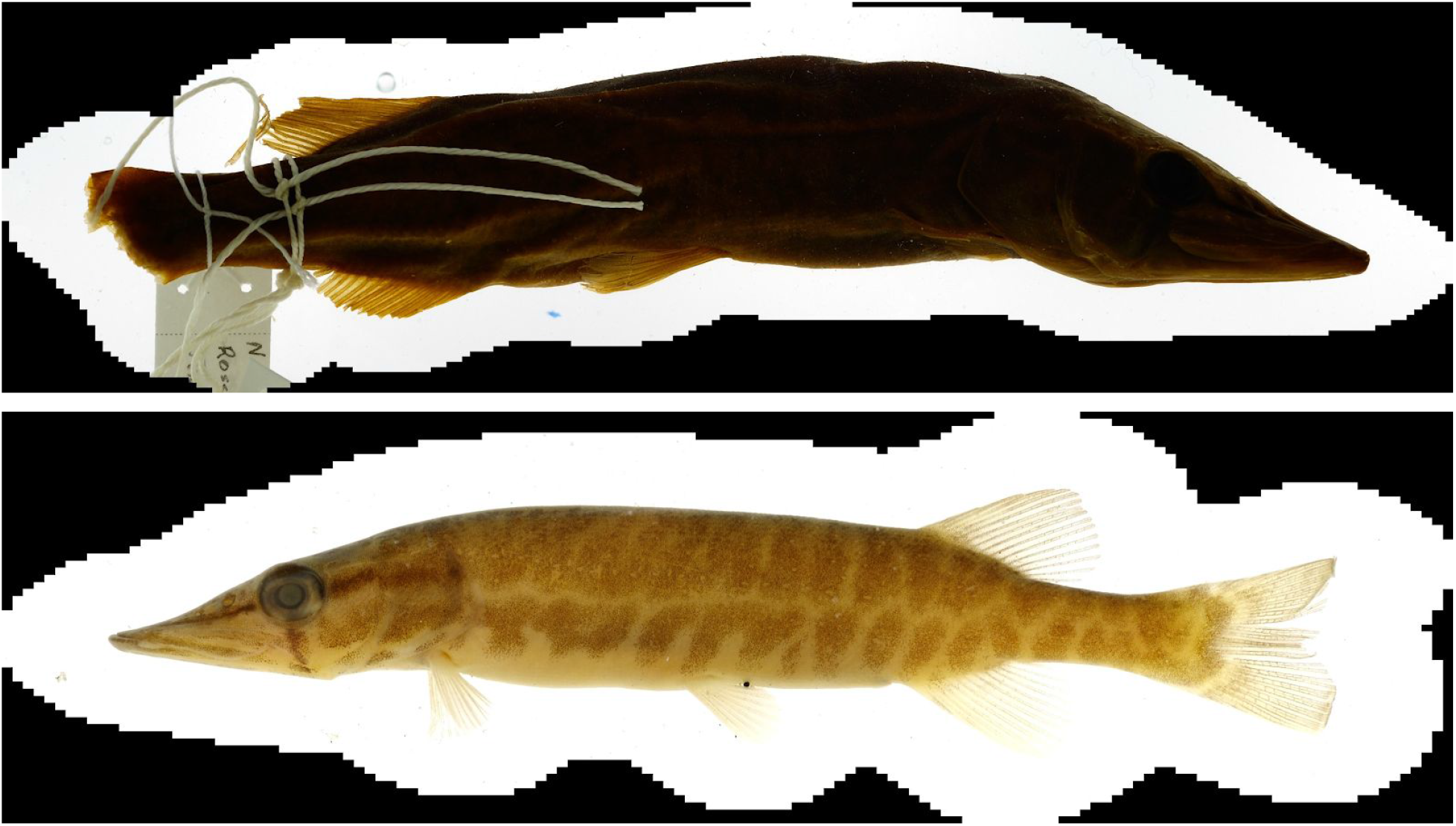
Examples of very low (1) and very high (10) quality images of Esox americanus

## 6. Results

### Low/High Subset Comparison

A t-test of F1 scores generated by several runs on the small balanced high and low quality subsets showed a small but significant difference in accuracy (0.41 vs 0.35, pval=0.031) (Figure 6).

**Fig. 6.**
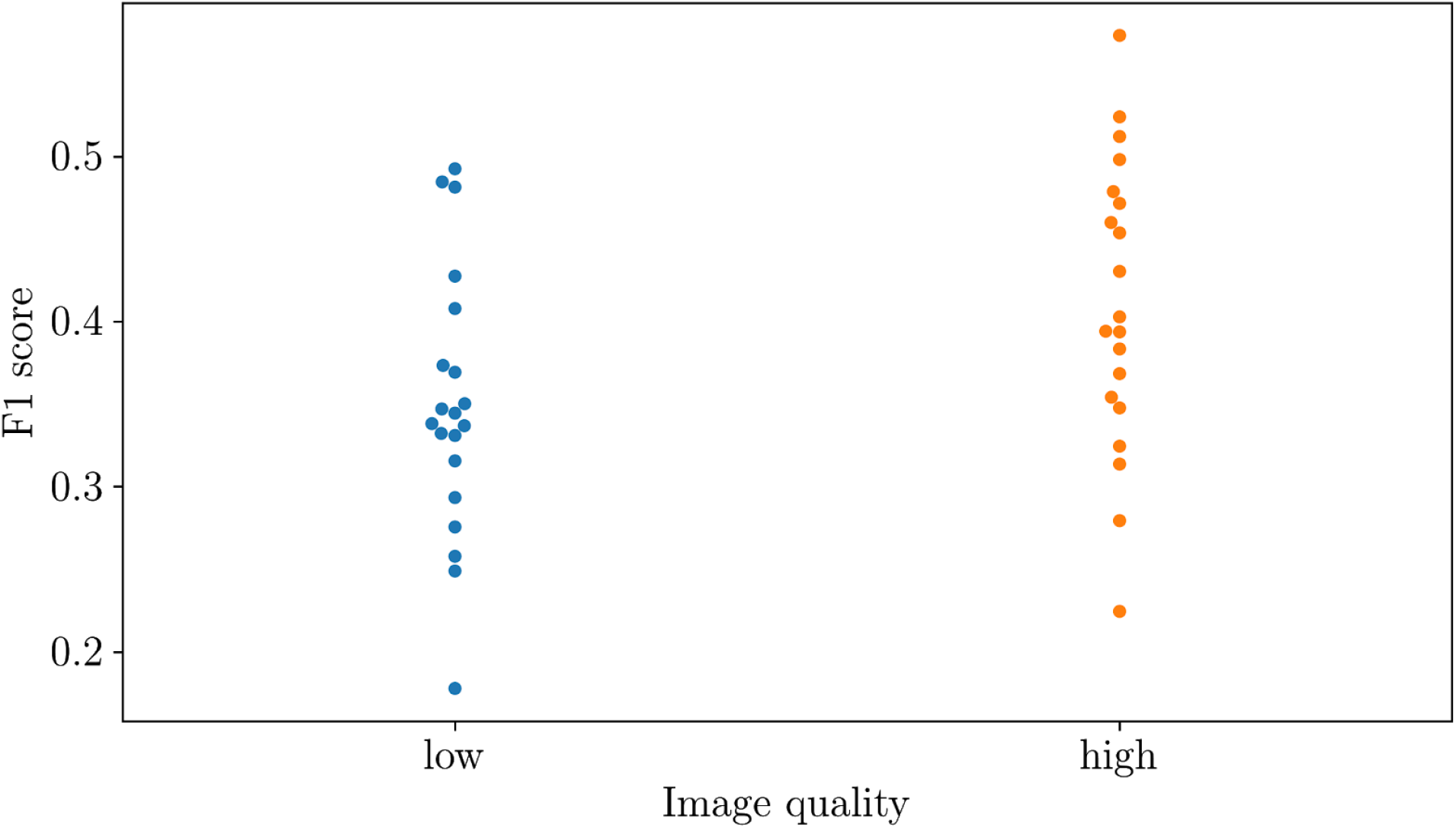
F1 test score across 19 trials on genus classification using low quality (mean 0.35) and high quality (mean 0.41). Using high quality images produced better F_1_ scores (0.41 vs 0.35, pval=0.031).

### Quality by Classification Outcome

Here we compared correctly classified vs misclassified images using a test set of 341 images (278 correctly classified and 63 misclassified). A confusion matrix (Figure 7) shows that most misclassifications occur between species within the same genus, although these misclassifications are not symmetric (e.g. 53% of *Notropis blennius* images were misclassified as *Notropis atherinoides*, while no *Notropis atherinoides* were misclassified as *Notropis blennius*).

**Figure 7.**
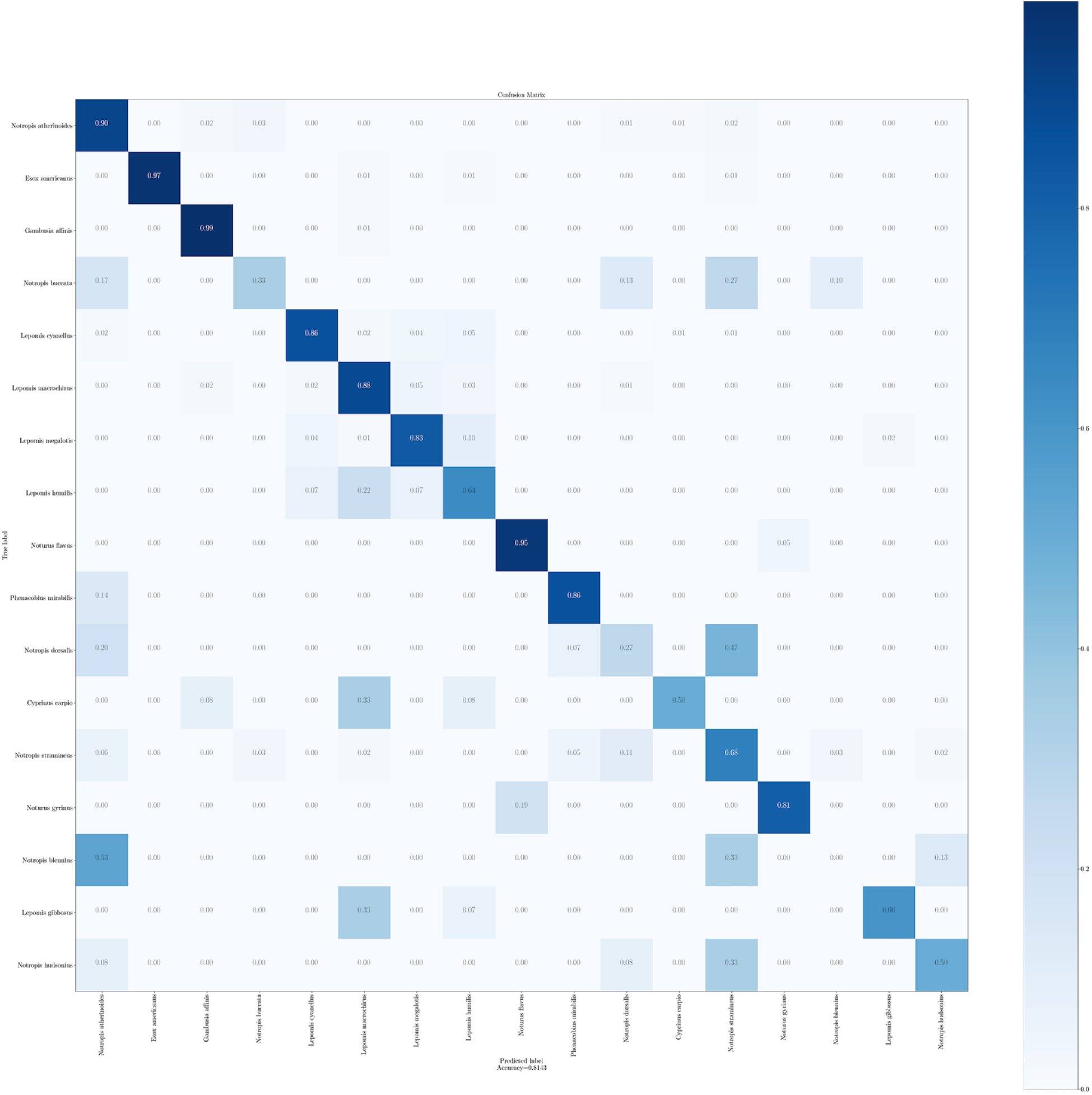
A confusion matrix of 341 species classifications. The diagonal represents correct classifications. The y-axis represents true labels, and x-axis are predictions.

Comparing the means of quality scores between correctly classified images reveals five quality features correlated with classification accuracy: if_curved (0.0179 vs 0.0634), if_parts_visible (0.8669 vs 0.8413), if_overlapping (0.1043 vs 0.1270), and image_quality (8.230 vs 8.079), and two negatively correlated: if_background_uniform (0.6151 vs 0.6984), and if_fins_folded (0.046512 vs. 0.025641). While image_quality is the strongest variable, a logistic regression which includes all features except image_quality (to avoid collinearity), reveals if_parts_visible (p-val = 0.0001) as the sole significant covariate (Table 4).

**Table 4:**
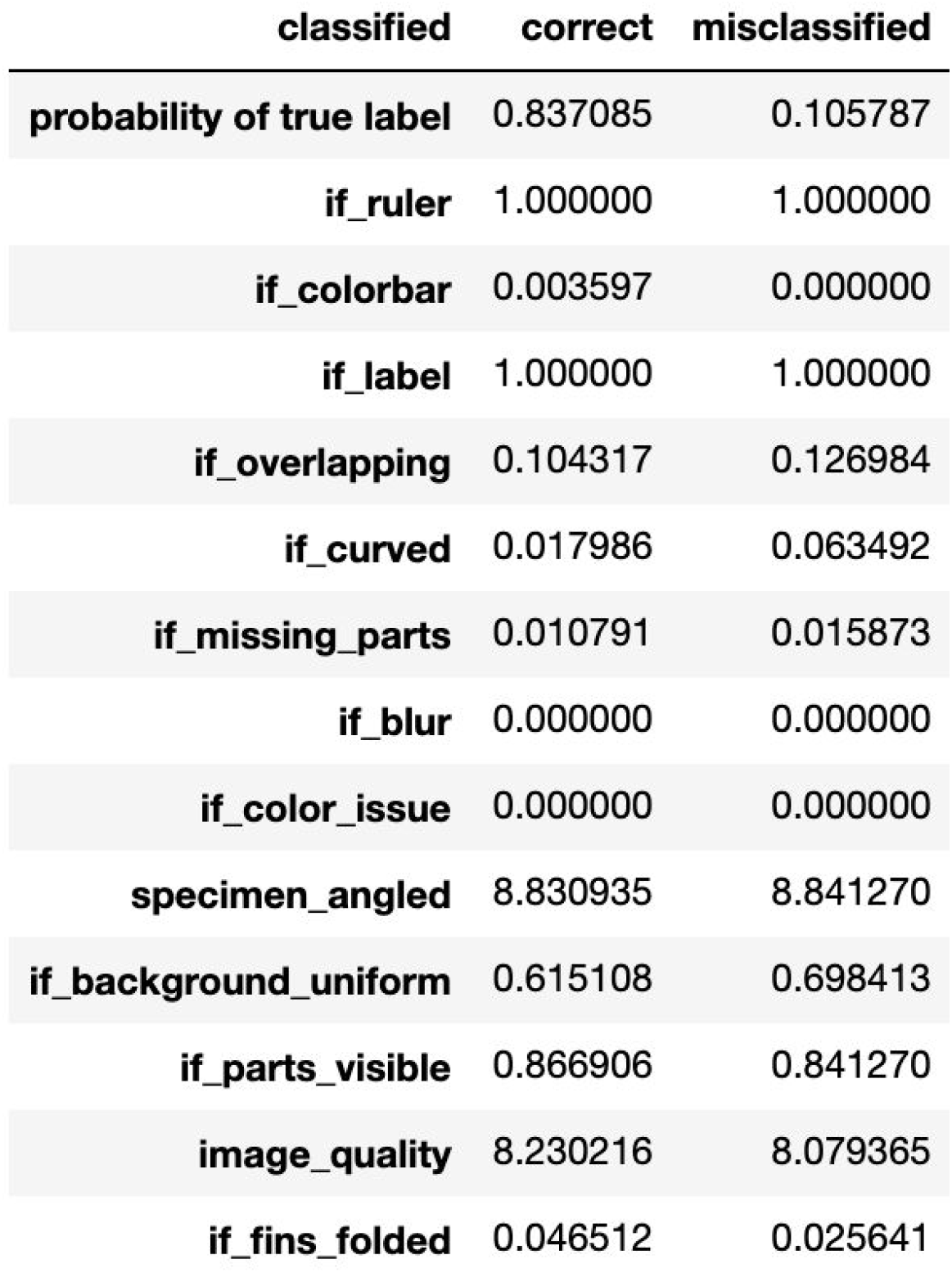
Mean of quality features by correctly classified and misclassified images

## 7. Discussion

In this paper we show that image quality measures do affect classification accuracy. This was demonstrated using two approaches – a dichotomous split of the image corpus using the manually-annotated image_quality metric, and a comparison of correctly and incorrectly classified images from the entire quality-annotated data set. Our abilities to discern the importance of quality are hampered by three factors: 1) a relative paucity of low-quality images in our dataset, 2) the nature of classification – some fish are simply more similar to their brethren – but we have attempted to control for this where possible, and 3) some taxa are inherently more difficult to position or illuminate for photography. Our results lead to a number of recommendations for assessing quality of images from biodiversity image repositories to support machine learning analyses.

The quality_score assigned by curators, while based on a rubric, does lend itself to some inter-rater error. We surmised that a composite metric of the binary quality items (e.g. if_curved, if_fins_folded, etc..) could represent a more objective score, and explored this, but it ultimately did not prove substantially better than “image_quality”.

The quality scores generated by our curators included some that are strictly technical (blur, color issues), those that would apply to any biodiversity catalog (“if_parts_visible”) and those that are specific to fish (e.g. “if_fins_folded”). We contend that all three types of quality (technical, biospecimen, taxon-specific) are important to include for biorepositories. The automated measurement of technical image quality, and possible higher-level judgments, can help accelerate the collection of this metadata. Metadata librarians may also find it useful to distinguish local and global image quality characteristics <u>^23,24^</u> from semantic quality features. Local features might include fin-level textures that would indicate lepidotrichia, global features such as large segmented areas as well as basic image characteristics such as color and shape. These characteristics are logically distinct from semantic quality judgments made by the curators in this project (“folded fins”/”label obstruction”), though automated semantic quality annotations are within the capabilities of neural networks.

Another metadata property that we have mentioned tangentially is provenance, particularly because of the batch effects introduced by disparate labs collecting and photographing specimens with different settings and equipment. Batch effects are a huge issue in all repositories and data coordinating centers, and due to the geographically localized nature of certain species and associated labs, this can sometimes present a “complete separation” problem, where controlling for the random effects of collection centers becomes impossible. This would suggest each center be encouraged to collect certain common species to serve as controls for normalization. Both provenance and quality are essential for large image repositories to address bias and confounds for downstream analyses.

Overall, observed that machine learning is hampered by its dependence on classification accuracy instead of more direct intermediate measures, for example, the number of features detected. Though this is an active area of research within the deep learning community, species detection will continue to present some confounds because of inherent heterogeneity discussed above. Certainly, one consideration specimen curators should be aware of is whether the intended use of the biorepository images is to classify real-world specimens. Certain types of low-quality images may serve to augment robustness in computer vision – a technique called “noise injection”. These uses suggest annotating for quality, rather than simply culling, is a preferable strategy. Quality metadata to aid robustness and generalizability in machine learning, rather than a narrow focus on pristine specimens, is an open area for future work.

Annotation of fish images is an ongoing project with BGNN, and a larger cohort of quality-curated images should validate the findings in this paper. With a larger data set we should also be able to conduct a cross-validation of low-high quality models against their complementary data set, to examine the effect of robustness training.

## 8. Conclusion

The main objective of this research was to determine if annotated image quality metadata impacted generic and species-level classification accuracy of a convolutional neural network. We conducted an empirical analysis to examine which specific quality annotations were most important for classification accuracy. We worked with a set of 22 metadata properties developed at Tulane University to evaluate the quality of large caches of 2D fish specimen images. Our key finding was that images with high-quality metadata metrics had significantly higher classification accuracy in convolutional neural network analysis than images with low-quality metadata metrics.

We offer a number of recommendations for assessing the quality of images from biodiversity image repositories to support machine learning analyses. This investigation serves as a baseline study of useful metadata for assessing image quality. The methodology and our approach also serves to inform other research that may examine image quality and impact on classification for other fishes, other specimens, and even other disciplines where the image is a central object. Overall the research conducted serves the needs of the BGNN project, and biologically-focused, machine-learning projects generally, for determining whether images for biodiversity specimen image repositories are useful for higher-level analysis.

## 9. Acknowledgments

We thank Tulane University Technology Services for assistance setting up servers for hosting images and sharing them with other BGNN investigators. We thank Justin Mann, Aaron Kern, Jake Blancher for help with refining the metadata form and gathering the metadata used in the project. The fish specimen images used in the project were obtained from Chris Taylor, Curator of Fishes and Crustaceans at the Illinois Natural History Survey (INHS). INHS is one of six fish collections participating in the Great Lakes Invasives Network (GLIN), five of which shared fish specimens images for use in the BGNN project.

## 10. Data availability

Raw data is available at bgnn.org/INHS. Reproducible code is available at https://github.com/hdr-bgnn/iqm

## 11. Acknowledgements

Research supported by NSF OAC #1940233 and #1940322.We acknowledge Tulane University Technology Services for server set-up and access supporting this research; Justin Mann, Aaron Kern, Jake Blancher for curation work; and Chris Taylor, Curator of Fishes and Crustaceans, INHS, for providing the fish specimen images and metadata (INHS support: NSF Award #1400769).

## Appendix

**Figure 8:**
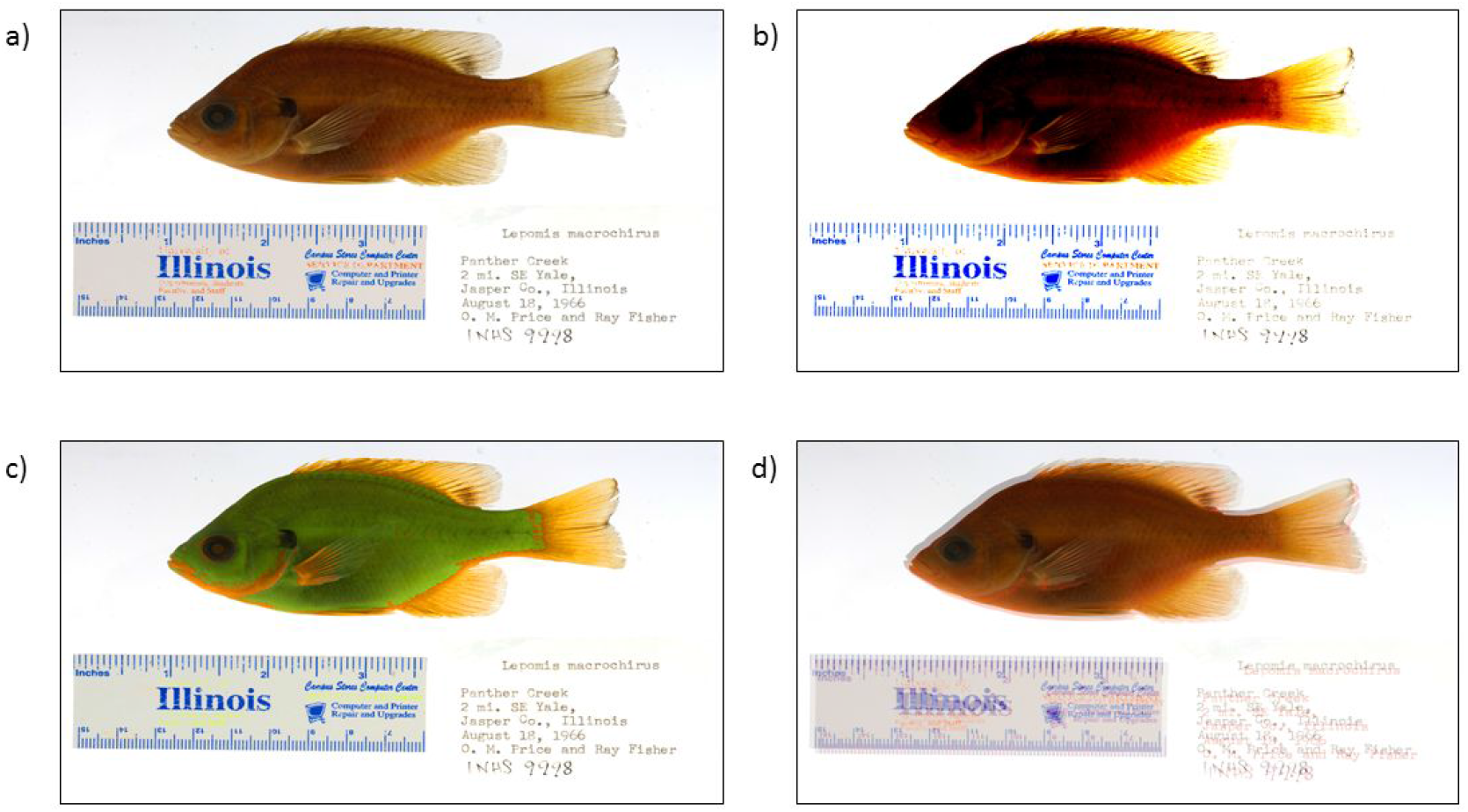
An example question in the questionnaire for color issues; a) original image, b) contrast issue, c) saturation issue, d) shifted colors.

